# Nanoscale Protonation Limits and Charge Density in Polymer Films Govern the Activity of Immobilized LacZ under Acid Stress

**DOI:** 10.64898/2026.01.28.702428

**Authors:** Huida Duan, Junxing Chen, Felicia Fianu, Wei Sun, Yifan Cheng

**Author notes:** Correspondence: Yifan Cheng, Wei Sun. Equal contributions (Huida Duan, Junxing Chen).

## Abstract

Under acidic conditions, polycationic polymer coatings function as protective immobilization supports through protonation-mediated local pH buffering. However, it remains unclear how polymer support design parameters, such as film thickness and charge density, govern that vital protonation process. Leveraging the precise control of film thickness and copolymer composition enabled by initiated chemical vapor deposition (iCVD), we systematically investigated how these parameters govern the protonation behavior of poly[glycidyl methacrylate-*co*-2-(dimethylamino)ethyl methacrylate] (pGD) thin films and, in turn, the activity of immobilized β-galactosidase (LacZ). Infrared spectroscopy suggests that proton penetration was capped at a depth of ∼250 nm in pGD with 65% DMAEMA, limiting the polycationic thickness in pGD films thicker than this value. Consistent with this limit, immobilized LacZ activity under acidic stress (pH 4) increased with protonated thickness up to ∼250 nm and then plateaued. Raising the polycationic monomer content from 25 to 65 mol% increased LacZ activity at pH 4 by up to 83%, consistent with a higher positive charge density providing stronger local pH buffering. To test whether this behavior depends on immobilization sites, we evaluated two approaches: random immobilization (via amine-epoxy ring-opening reactions) and site-directed immobilization (via SpyCatcher/SpyTag binding). Directed immobilization preserved higher LacZ activity than random immobilization, but the protonation-dependent protection trend remained consistent for both strategies. These findings establish protonation depth and charge density as tunable design parameters for polycationic immobilization supports that stabilize enzymes under acidic conditions.

## 1. Introduction

Immobilized enzymes are foundational tools in biotechnology and industrial biocatalysis, offering advantages such as enhanced stability, recoverability, and recyclability.^1– 3^ Despite these advantages, maintaining the catalytic efficiency of immobilized enzymes under suboptimal environmental conditions, particularly at low pH, remains a critical challenge.^4,5^ The support material’s microenvironment, especially its surface chemistry and morphology, plays a critical role in preserving enzyme activity by regulating local pH, hydrophilicity and conformational stability.^6,7^ Functional polymer coatings have emerged as versatile platforms that allow functional groups to be tailored for robust enzyme attachment, including covalent anchoring, and microenvironment modulation.^8,9^ Among these, epoxide-functionalized polymers are widely adopted for enzyme immobilization because they form stable covalent linkages with nucleophiles, such as the amines in amino acid side chains on enzyme surfaces.^10^

Meanwhile, high pH-responsive polymers, particularly those bearing tertiary amine groups such as poly[2-(dimethylamino)ethyl methacrylate] (pDMAEMA) have shown promise in buffering local microenvironments by changing their ionization state in response to bulk pH shifts.^11–13^ For instance, pDMAEMA brushes grafted onto surfaces underwent reversible transition between collapsed (neutral) and swollen (protonated) conformations as pH changes, thereby modulating enzyme accessibility and stability.^13,14^ These materials are also advantageous due to their tunable charge density, which can be adjusted by controlling the fraction of DMAEMA in copolymer formulations.^15–17^ Together, these advancements suggest a path toward designing enzyme immobilization platforms that not only anchor enzymes but also tailor their microenvironment, particularly under pH extremes.

Despite the promise of pH-responsive polymers for enzyme immobilization, a fundamental understanding of how polymer thickness and charge density modulate protonation behavior—and how this, in turn, influences biocatalytic performance— remains lacking.^18,19^ In addition, while both random immobilization (e.g., amine-epoxy ring opening) and (site-)directed immobilization (e.g., SpyTag/SpyCatcher binding) strategies are established,^20^ studies integrating site-specific orientation with charge-tunable polymer and evaluating performance under acidic conditions remain limited. Clarifying these relationships would enable more effective local pH modulation and improved activity retention of immobilized enzymes under acidic conditions. Moreover, identifying the minimum polycationic-support thickness required for enzyme protection would guide cost-effective support design.^19,20^

In this study, we derived design rules for pH-responsive polymer supports by independently tuning film thickness and positive charge density. We copolymerized glycidyl methacrylate (GMA, “G”) and 2-(dimethylamino)ethyl methacrylate (DMAEMA, “D”) to form pGMA-*co*-DMAEMA (pGD) thin films, where G supplies epoxide groups for covalent protein conjugation^21,22^ and D supplies tertiary amines that protonate under pH < pKa (∼8.4) to generate positive charge.^7^ To precisely control thickness and copolymer composition, we synthesized these supports using initiated chemical vapor deposition (iCVD), an all-dry, solvent-free vapor-deposition polymerization method^23,24^. During the iCVD process, monomer and initiator vapors (typically tert-butyl peroxide, TBPO, as the initiator) are metered into a reactor held at moderate vacuum (commonly ∼0.1–1 Torr).^25^ Monomers adsorb on a cooled substrate (often ∼20–40 °C), while TBPO thermally decomposes to generate radicals that initiate surface-confined chain-growth polymerization.^26^ As a result, iCVD enables nanometer-scale thickness control and high retention of monomer functional groups (e.g., epoxides and tertiary amines) under solvent-free, mild-temperature conditions.

In addition to iCVD-synthesized polycationic supports, we incorporated self-assembled monolayers (SAMs) bearing epoxysilane and tertiaryaminosilane to access the ultrathin limit of a single molecular layer (typically ∼0.5–3 nm, depending on chain length and substrate).^27,28^ SAMs are a well-established, solution-phase surface modification approach for presenting reactive groups and mediating covalent enzyme immobilization,^27,29^ and here they provide an extreme-thickness benchmark for comparison with tens-hundreds of nm iCVD films.

Combining iCVD films and silane-based SAMs, we fabricated chemically parallel polycationic immobilization supports with well-defined thicknesses spanning ∼1 to 800 nm to systematically examine protonation behavior and enzyme activity at pH 4. To probe the interplay between film thickness and charge density, we varied the DMAEMA content in p(GMA-co-DMAEMA) films (25% and 65%). We hypothesized that protonated tertiary amines establish an electrostatic barrier that limits further proton penetration, leading to a self-limiting protonation regime once the film exceeds a critical thickness. This effect, in turn, may modulate local pH buffering and the protection of immobilized enzymes under acidic stress. Overall, our findings reveal protonation depth and charge density as tunable design parameters for engineering polycationic immobilization supports that stabilize enzymes in low-pH environments.

## 2. Materials and Methods

### 2.1 Nomenclature

In this study, we employed four monomers: glycidyl methacrylate (GMA) and N,N-dimethylaminoethyl methacrylate (DMAEMA) for iCVD, and 3-glycidylpropyl trimethoxysilane (GPTMS) and 3- (dimethylamino)propyltrimethoxysilane (DMAPTMS) for SAM coatings. We adopted a concise naming convention in which “G” denotes an epoxide-bearing monomer (GMA or GPTMS) and “D” a tertiary amine monomer (DMAEMA or DMAPTMS), while a leading lowercase “p” signifies an iCVD synthesized thin film and a leading uppercase “S” signifies a SAM monolayer; thus, for example, pGD refers to an iCVD film copolymerized from GMA and DMAEMA and SGD to a SAM composed of GPTMS and DMAPTMS.

### 2.2 Chemical reagents

All reagents were used as received: the iCVD monomers glycidyl methacrylate (GMA, 97.0% purity with ∼0.01% hydroquinone monomethyl ether stabilizer), N,N-dimethylaminoethyl methacrylate (DMAEMA, 98% purity with 700–1000 ppm monomethyl ether hydroquinone inhibitor) and the initiator tert-butyl peroxide (TBPO, 98% purity) were obtained from Sigma-Aldrich Inc. (St. Louis, MO, USA), while the SAM reagents 3-glycidylpropyl trimethoxysilane (GPTMS, 98%) and 3- (dimethylamino)propyl trimethoxysilane (DMAPTMS, 96%) were procured from TCI America via Fisher Scientific Company LLC (Suwanee, GA, USA).

### 2.3 Preparation of iCVD-Derived Polycationic Nanolayer Supports

Polymers were deposited via iCVD to serve as LacZ immobilization supports. Glycidyl methacrylate (GMA) introduced epoxy groups for covalent enzyme attachment and N,N-dimethylaminoethyl methacrylate (DMAEMA) that impart pH-responsive buffering capacity to the coating. Tert-butyl peroxide (TBPO) was used as the radical initiator, in line with established iCVD polymerization processes.^30^ All precursors were vaporized and metered into the reactor at a combined flow of 3–5 sccm, with TBPO held constant at 0.6 sccm. The substrate stage was maintained at 35°C under a total chamber pressure of 0.6 Torr. A resistively heated filament array at 220°C generated the free radicals necessary to initiate polymerization. Corning™ polystyrene 96-well plates were mounted on a custom aluminum platen for efficient thermal coupling to the reactor’s cooled stage, and a silicon slide atop the wells allowed in situ thickness monitoring via a He–Ne interferometer (350–700 nm; Thorlabs Inc., Newton, NJ, USA). Deposition parameters yielded 12–200 nm coatings on the Si slide (stage temperature ∼65°C, P_m_/P_sat_ ≈ 0.01). Considering the quadratic dependence of deposition rate on monomer partial pressure at low P_m_/P_sat_ (<0.04 for acrylates), corresponding films deposited in 96-well plates where monomer accumulation increases local P_m_, are estimated to be 50–800 nm. Detailed gas flow, P_m_/P_sat_ ratios, deposition rates and copolymer composition are compiled in **Tables S1, S2 and S3**.

### 2.4 Fabrication of Self-Assembled Monolayers (SAMs)

To simulate critical functional chemistries of our iCVD films, we formed mixed SAMs from 3-glycidyloxypropyltrimethoxysilane (GPTMS) and 3- (dimethylamino)propyltrimethoxysilane (DMAPTMS) on both 96-well plates and silicon wafers. Substrates were first activated by oxygen plasma (45 W, 500–900 mTorr, 3 min; Harrick Plasma Inc., NY, USA) to generate surface hydroxyl groups for silane coupling. Separately, 1% v/v ethanolic solutions of GPTMS and DMAPTMS were stirred at 650 rpm and 20°C for 1 h, then combined to yield a mixture containing 35% GPTMS and 65% DMAPTMS— matching the DMAEMA fraction in the pGD(65) iCVD film. Plasma-treated substrates were submerged in the GPTMS/DMAPTMS blend for 18 h at room temperature to allow silanization. Finally, SAM-coated surfaces were rinsed sequentially with ethanol and deionized water and dried under ambient conditions, following established processes.^30^

### 2.5 Ellipsometry

To confirm the iCVD film thickness prior to LacZ immobilization, ex-situ measurements were performed on silicon wafer samples using a J.A. Woollam alpha-SE spectroscopic ellipsometer (Lincoln, NE, USA). Data were collected at three incident angles (65°, 70° and 75°) and fit to a Cauchy–Urbach model to extract thickness values. Each sample was measured at three separate locations, and the average thickness was reported to ensure consistency before proceeding with enzyme immobilization and downstream characterizations.

### 2.6 Fourier Transform Infrared spectroscopy

Fourier-transform infrared spectroscopy (FTIR) analysis was conducted to verify the presence and preservation of characteristic functional groups in the polymer films synthesized by iCVD. The FTIR spectroscopy of all immobilization supports was performed utilizing a Thermo Scientific Nicolet iS50 Model (Austin, TX, USA). To characterize the composition of copolymers, FTIR analysis was conducted in transmission mode. The experiment employed attenuated total reflectance Fourier transform infrared spectroscopy (ATR-FTIR) using the Thermo Scientific Nicolet iS50 Model, which was equipped with a VariGATR grazing angle ATR accessory (GAT-V-N18, Harrick Scientific Products, NY). The incident angle of the ATR accessory, 60°, was utilized in this study. A MCT detector cooled with liquid nitrogen was utilized over the spectral range of 600-4000 cm^-1^, with a resolution of 4 cm^-1^. Measurements were averaged over 128 scans to improve the signal-to-noise ratio. Baseline correction was applied by subtracting a background spectrum obtained from an uncoated Si wafer substrate.

### 2.7 Characterization of polymer protonation

The protonation behavior of synthesized polymer films (pGD and SGD) was examined by ATR-FTIR spectroscopy. Specifically, polymer-coated substrates were immersed in acidic solution (pH 4, HCl) or basic solution (pH 10, NaOH) for 5 min, followed by air-drying for 1 min. FTIR spectra of polymer films before and after soaking were collected and compared to confirm the occurrence of protonation. Furthermore, to quantify the protonation extent across polymer films of varying thicknesses, FTIR spectra were subsequently collected in transmission mode. The area ratio of the newly emerged N–H bending vibration peak (indicative of protonation) to the characteristic carbonyl (C=O) peak was calculated.

### 2.8 Bioengineering of enzymes

#### 2.8.1 Construction of pBAD-LacZ, pBAD-LacZ-spyTag, and pBAD-SpyCatcher Expression Plasmids

The wild-type LacZ expression plasmid (pBAD-LacZ-WT) was constructed by amplifying the LacZ gene (UniProt ID: P00722) via colony PCR using the primer pair LacZ-WT-NdeI-F and LacZ-WT-6xHis-R. The resulting PCR product was digested with NdeI and HindIII and ligated into a pBAD vector backbone pretreated with the same restriction enzymes. To construct the pBAD-LacZ-spyTag plasmid, a SpyTag sequence fused to a glycine-serine (GS) linker was PCR-amplified using the primers spyTag-GSlinker-F and spyTag-GSlinker-R, and recombined with a LacZ-WT PCR product amplified with pBAD-LacZ-WT-spy-F and pBAD-LacZ-WT-spy-R, yielding a construct with the SpyTag at the N-terminus of LacZ. For the pBAD-SpyCatcher (pBAD-SC) plasmid, the SpyCatcher coding sequence (GenBank: AFD50637.1) was codon-optimized and synthesized by IDT. The gene was amplified using SC-NdeI-F and SC-HindIII-6xHis-R, digested with NdeI and HindIII, and ligated into a pBAD vector treated with the same enzymes. Sequences of all primers used in these constructions are provided in **Table S**4.

#### 2.8.2 Expression of LacZ and SpyCatcher

To express wild-type LacZ (LacZ-WT), LacZ-SpyTag (LacZ-ST), and SpyCatcher (SC) proteins, the plasmids pBAD-LacZ, pBAD-LacZ-SpyTag, and pBAD-SpyCatcher were individually transformed into chemically competent *E. coli* DH10B cells. Transformants were selected on LB agar plates containing 100 µg/mL ampicillin (LB-Amp100) and incubated overnight at 37°C. A single colony from each plate was used to inoculate 5 mL of LB-Amp100 medium and cultured overnight at 37°C. Then, 1 mL of this preculture was transferred into 5 L of fresh LB-Amp100 medium. The cultures were shaken at 37°C until the optical density at 600 nm (OD_600_) reached 0.4–0.6, and protein expression was induced by adding 0.2% (w/v) L-arabinose. For LacZ-WT and LacZ-ST, induction was continued for 24 h at 37°C, while for SC the culture was maintained for 24 h at 18°C to promote soluble expression. Cells were harvested by centrifugation at 7000 × g for 15 min at 4°C, and the resulting cell pellets were stored at −80°C until use.

#### 2.8.3 Purification of His-Tagged Proteins via Ni-NTA Chromatography

Cell pellets were thawed and resuspended in 14 mL lysis buffer (20 mM Tris-HCl, pH 8.8, 400 mM NaCl, 20 mM imidazole, protease inhibitors), incubated at 4 °C for 30 min, and lysed by sonication (Branson Sonifier 450, 50% output, pulse mode: 1 s on/off, total duration 15 min, with 1-min pauses every 5 min) in an ice-water bath. Lysates were clarified by centrifugation (16,000 × g, 4°C, 30 min), and the supernatants incubated with 5 mL pre-equilibrated HisPur™ Ni-NTA resin (Thermo Scientific) for 1 h at 4°C with continuous rotation. The resin slurry was transferred onto a Poly-Prep® chromatography column, washed three times with 20 mL wash buffer (20 mM Tris-HCl, pH 8.8, 400 mM NaCl, 20 mM imidazole, 2 mM DTT), and proteins eluted in five fractions of 3 mL each using elution buffer (20 mM Tris-HCl, pH 8.8, 400 mM NaCl, 500 mM imidazole, 2 mM DTT). Eluates were concentrated via Amicon Ultra-15 centrifugal filters (30 kDa cutoff for LacZ variants; 3 kDa for SC), buffer-exchanged into storage buffer (20 mM Tris-HCl, pH 8.8, 400 mM NaCl, 2 mM DTT, 10% glycerol) using Cytiva PD-10 Sephadex G-25M desalting columns, flash-frozen in liquid nitrogen, and stored at −80°C.

### 2.9 Enzyme immobilization

For RI, LacZ-WT solution (200 µL, 0.06 mg/mL) was added to a Corning 96-well plate previously coated with polymer layer. The plate was incubated statically at 37 °C for 24 h to achieve covalent attachment via amine-epoxide coupling. DI proceeded in three sequential steps: (i) random immobilization of SC, (ii) blocking of unreacted epoxides with glycine, and (iii) site-specific capture of LacZ-ST via SC/ST interaction. Briefly, SC (200 µL, 0.05 mg/mL, 2.8 × 10^−4^ mM) was randomly immobilized under conditions identical to LacZ-WT. Remaining epoxide groups were blocked using 1.0 M glycine solution following Gao et al. Subsequently, LacZ-SpyTag solution (0.06 mg/mL, 1.0 × 10^−4^ mM) was added, and the plate was incubated at 25 °C with shaking (30 rpm) for 2 h as previously described. After immobilization, residual enzyme solutions were removed, wells were rinsed twice with 1× PBS to remove unbound proteins, and immobilized LacZ was immediately subjected to activity assays and protein quantification.

### 2.10 Protein Quantification and Enzyme Activity Assays

Protein quantification of immobilized LacZ variants was performed using a Pierce bicinchoninic acid (BCA) assay kit (Thermo Fisher Scientific, Waltham, MA, USA). Briefly, 25 μL of 0.1 M phosphate buffer (pH 7.0) was added to each enzyme-immobilized well, followed by 200 μL of BCA working reagent. Plates were mixed thoroughly for 30 s on a plate shaker and incubated at 37 °C for 30 min. Protein quantification was performed for all immobilized proteins, including randomly immobilized LacZ-WT, and sequentially for SC and LacZ-ST in directed immobilization. Immobilization yield (IY) was calculated using the formula:

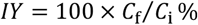

where *C*_i_ represents the initial enzyme concentration before immobilization, and *C*_f_ is the concentration equivalent of immobilized enzyme determined by the BCA assay.

Enzyme activity assays of immobilized and free LacZ were carried out at 37 °C using o-nitrophenyl-β-D-galactopyranoside (ONPG) as substrate. Activity was measured across a pH range using 0.1 M lactate buffers (pH 4) and 0.1 M phosphate buffers (pH 7). Prior to the assays, polymer-coated plates with immobilized LacZ were equilibrated at 37 °C for 10-20 min. Reactions were initiated by adding ONPG solutions at concentrations of 0 and 16.6 mM. The generation of o-nitrophenol was monitored at 420 nm for 5 min using a BioTek Synergy H1 Multi-Mode microplate reader. The normalized initial rate 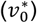 defined as the initial reaction rate measured at 16.6 mM ONPG normalized by the concentration of LacZ, was employed to compare enzyme activity among samples.

### 2.11 Enzyme Activity under Different Protonation Depths

To evaluate the influence of polymer protonation depth on the catalytic performance of immobilized enzymes, LacZ activity was measured for both RI and DI configurations on iCVD-synthesized pGD(65) films of different thicknesses (50, 100, 200, 400, 800 nm). The DI configuration served as a control to minimize orientation variability, ensuring that differences in activity reflect the microenvironmental effects induced by polymer protonation. Enzyme activity was determined at both neutral (pH 7) and acidic (pH 4) conditions. The initial reaction rate (*v*_0_). Activities were also compared against free enzyme and immobilization on fully aminated SAM surfaces (SGD(65)) to distinguish protonation-driven effects from those due to surface chemistry or total charge density alone. The results were used to establish correlations between enzyme activity and the effective protonated layer thickness obtained from FTIR measurements.

### 2.12 Statistical analysis

All experiments were performed in triplicate (n = 3). Data analysis and visualization were carried out using RStudio and OriginLab, respectively. Significant differences among treatments were assessed by analysis of variance (ANOVA), followed by Tukey’s Honest Significant Difference (HSD) post-hoc test to pinpoint specific differences between groups. Results were reported at a significance level of 0.05 and visualized using compact letter displays to summarize Tukey’s test outcomes.

## 3. Results and Discussion

### 3.1 Characterization of immobilization supports and protonation

To illustrate the acid-induced protonation of tertiary amine groups in both pGD and SAM systems (**Figure 1a**), we first characterized the chemical structures and molar compositions of the thin films synthesized by iCVD, along with the corresponding SAMs, through FTIR spectroscopy (**Figure 1b, c**). First, we focused on several characteristic absorption peaks associated with key functional groups relevant to enzyme immobilization and microenvironment modulation. The absorption peak at approximately 900 cm^−1^ corresponds to epoxide ring-deformation vibrations from the monomer glycidyl methacrylate (GMA) in pGD(65) and its SAM counterpart SGD(65).^31–33^ The presence of this peak confirms the retention of epoxide groups, which serve as critical bioconjugation sites for enzyme attachment. This mechanism is widely reported to facilitate multipoint attachment and enhanced enzyme stability and activity in immobilized systems.^34^

**Figure 1.**
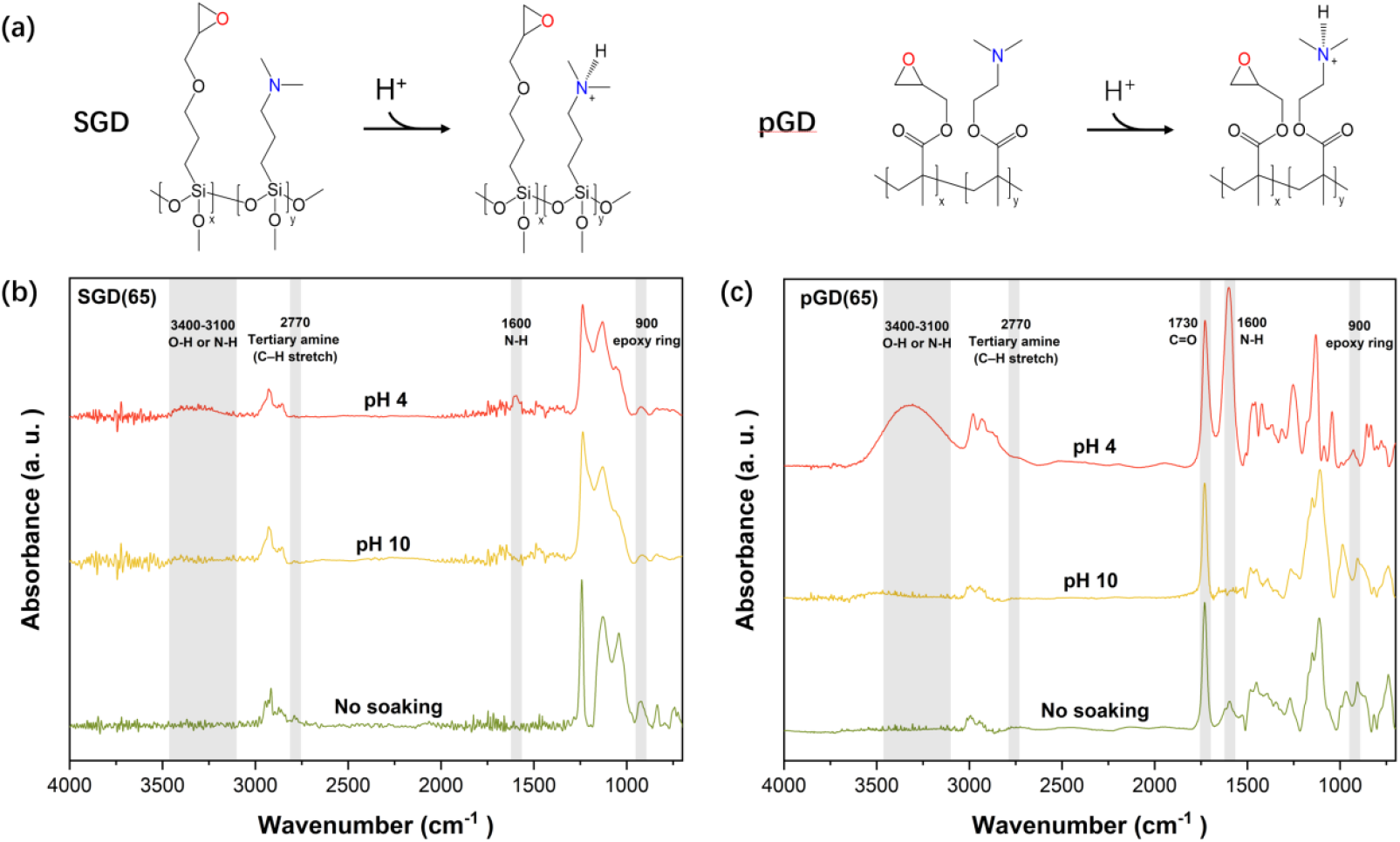
FTIR characterization of iCVD-synthesized thin films (pGD) and its self-assembled monolayer analog (SGD). (a) Reaction scheme showing protonation of the tertiary amine group in DMAEMA and DMAPTMS at acidic condition (pH 4). (b) FTIR spectra for SGD with 65%mol of D, SGD(65). (c) FTIR spectra for the iCVD-synthesized thin film with 65%mol of D, pGD(65).

Additionally, a weak absorption around 2770 cm^−1^ arises from symmetric C–H stretching vibrations of methyl groups linked to nitrogen atoms in tertiary amines. The consistent detection of this peak in both pGD(65) and SGD(65) indicates successful incorporation of tertiary amine groups derived from DMAEMA (iCVD) and DMAPTMS (SAMs).^7^ Furthermore, a strong carbonyl (C=O) stretching vibration at 1730 cm^−1^ was clearly observed in all iCVD-based polymers but absent in SAMs, confirming successful polymerization and correct structural formation. The disappearance of absorption peaks around the vinyl C=C stretch (∼1640 cm^−1^) further supports complete monomer conversion during iCVD synthesis.

To confirm protonation of the tertiary amine groups, spectra were compared before and after exposure to acidic (pH 4) and basic (pH 10) solutions. Acid treatment produced two new features: (i) a broad N–H stretch at 3000–3200 cm^−1^ and (ii) an N–H bending band near ∼1600 cm^−1^, both characteristic of protonated tertiary amines. These features were absent under basic conditions.^35^ Notably, these functional groups are derived from DMAEMA, whose tertiary amine has a known pKa of ∼8.4.^7^ In polymerized form (pDMAEMA), the effective pKa is somewhat lower (7.0–7.5) due to local electrostatic and steric effects.^36,37^ Thus, at pH 4, essentially full protonation is anticipated, in alignment with the observed FTIR features.^38^ In contrast, the absence of these bands in basic conditions indicates deprotonation and neutrality of tertiary amine groups.

The functional significance of these findings is substantial. Epoxide moieties ensure covalent enzyme binding, while tertiary amines confer pH-responsive charge modulation. Upon protonation at acidic conditions, the polymer/SAM surfaces become positively charged, creating a microenvironment capable of buffering local pH and protecting immobilized enzymes from external acidity. Literature documents on how buffers with ionizable groups close to enzymes help preserve activity under harsh conditions by maintaining local pH and shielding protein structure.^39^

In summary, FTIR analyses (**Figure 1b, c**) clearly verify the successful incorporation and preservation of epoxide functionalities essential for bioconjugation and ionizable tertiary amine groups crucial for pH-responsive modulation. The acid-induced protonation verified by FTIR further substantiates the pH-responsive capability of our designed polymeric supports. Collectively, these structural and functional confirmations underscore the suitability of our iCVD-synthesized polymer films and SAMs as advanced immobilization platforms.

### 3.2 Nanoscale Confinement of Proton Penetration

To further investigate how protonation varies with polycationic polymer thickness, pGD(65) films with varying thicknesses were immersed in an acidic solution (pH 4, HCl) for 5 minutes, then air-dried for 1 minute prior to analysis. The resulting FTIR spectra were analyzed (**Figure 2**). Specifically, protonation was tracked by the area ratio of the N–H bending vibration peak (∼1600 cm^−1^) relative to the carbonyl (C=O) peak (∼1730 cm^−1^). As shown in Figure 2, the protonation ratio increased with thickness up to ∼200 nm, after which it declined for thicker films. This observation suggests that protonation is a surface-limited process in thicker films, where protons predominantly react near the film interface, leaving the inner regions unprotonated within the experimental timeframe.

**Figure 2.**
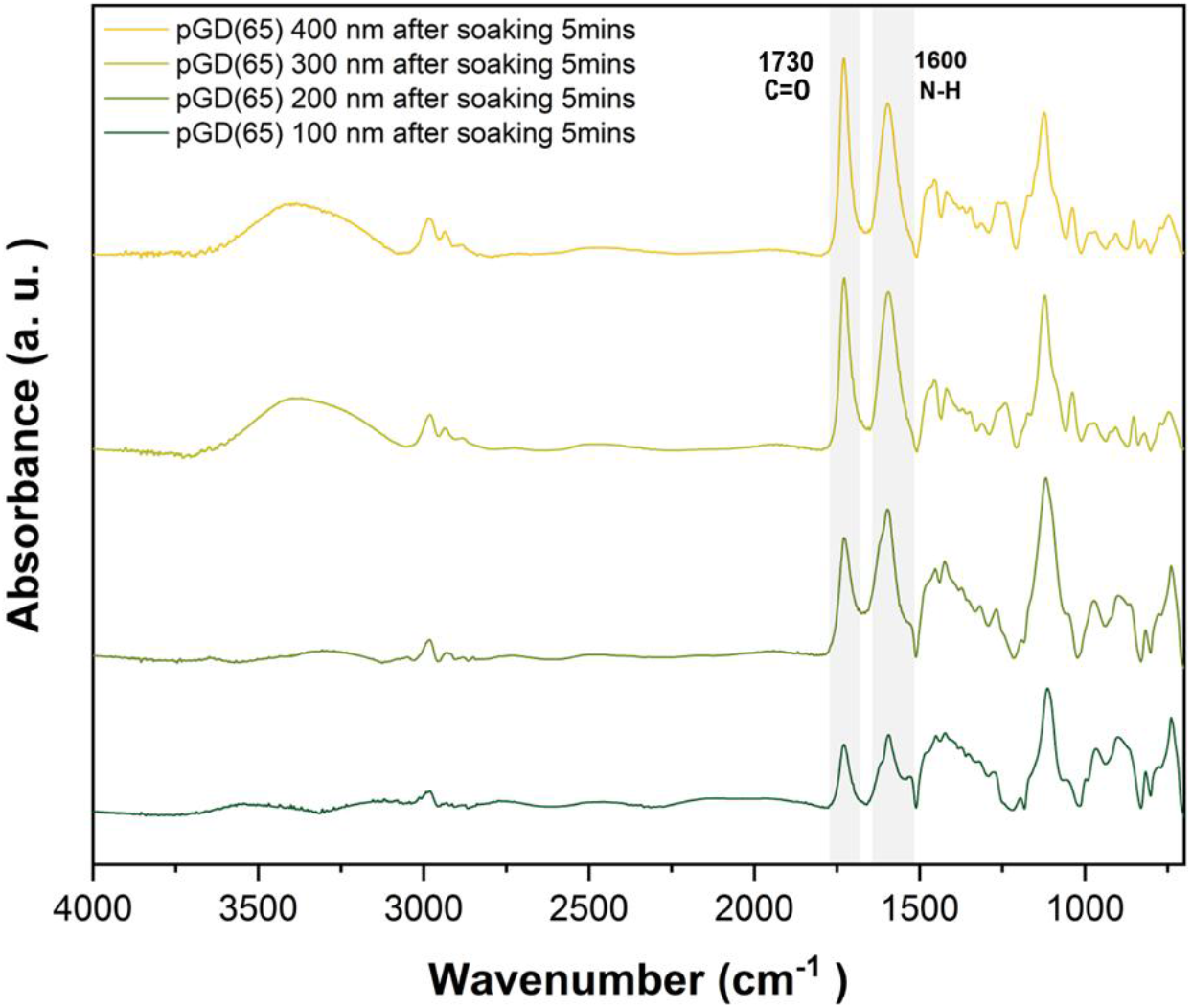
FTIR spectra of pGD(65) polymer films with different thicknesses after soaking in acidic solution (pH 4) for 5 min.

This thickness-dependent protonation behavior aligns with diffusion–reaction limited models observed in other polymer systems. Nguyen et al.^40^ demonstrated that strong-acid doping of conjugated polymer films results in surface-enriched dopant profiles, with a decaying concentration extending only tens of nanometers into the bulk, due to slower dopant diffusion into already charged surface layers. Similarly, Kolesov et al.^41^ reported that solution-phase doping of semiconducting polymers was confined to a thin interfacial layer, with penetration depths limited by both diffusion kinetics and local electrostatic interactions. In our case, as the pGD(65) film becomes protonated near the surface, the resulting positive charges attract counterions (Cl^−^) and increase osmotic swelling locally. This swelling can create a denser, gel-like skin that hinders further proton penetration into the unprotonated bulk. Another possible mechanism for this self-limiting behavior arises from the electrostatic repulsion exerted by positively charged, protonated tertiary amines, on the incoming protons. The formation of such a protonated surface barrier has been well-documented in pH-responsive polymer systems, where the interplay between ion diffusion, swelling, and fixed charge density results in a self-limiting penetration depth.^42^

For quantitative comparison, the N-H/C=O area ratio was calculated for each thickness and normalized by the fully protonated 100 nm polymer film. These normalized protonation percentages for films of varying thicknesses are presented in **Figure 3**. The results indicate that polymer films thinner than 200 nm can be fully protonated, whereas thick films showed partial protonation, suggesting the maximum protonation depth lies between 200 and 300 nm. The estimated protonated layer thickness was calculated by multiplying the as-synthesized iCVD film thickness, determined by ellipsometry in ambiance, by the protonation degree (%) derived from FTIR spectra. The resultant estimated protonation depth plateaued ∼250 nm, which we term the maximum proton penetration depth 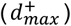, beyond which additional thickness no longer contributes to the protonated volume. Here we did not consider the swelling of the film in aqueous conditions.

**Figure 3.**
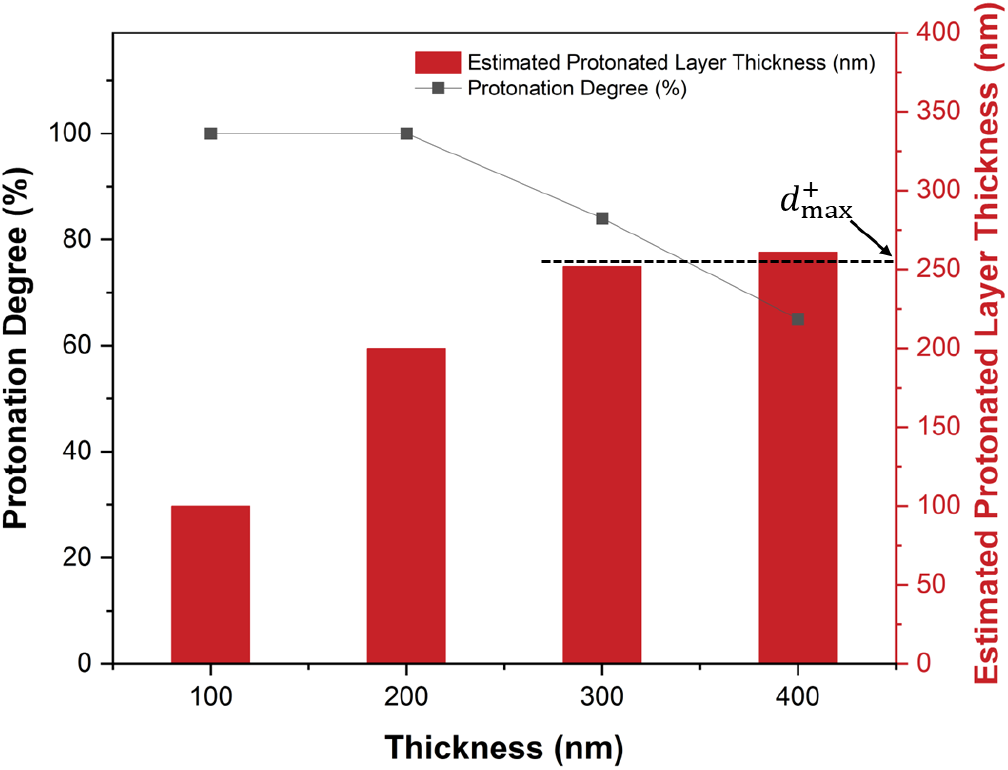
Normalized protonation degree (%) and estimated protonated layer thickness calculated from FTIR peak area ratios.

Furthermore, theoretical studies on weak polyelectrolyte brushes indicate that intra-chain charge repulsion and Donnan equilibrium effects can significantly hinder further ion uptake into dense polymer phases, unless external stimuli (e.g., increased ionic strength or prolonged exposure time) are applied.^43,44^ In our pGD(65) films, the protonation depth was not enhanced by simply increasing thickness, suggesting that the interior regions are effectively shielded from proton diffusion under ambient conditions.

### 3.3 Effect of Protonated Layer Thickness on Immobilized Enzyme Activity

Building upon the FTIR and quantitative analyses described above, which revealed a protonation depth of ∼250 nm under acidic conditions, we next investigated how this protonation depth influences the catalytic performance of immobilized LacZ. **Figure 4a, b** show the catalytic activities of LacZ immobilized via random immobilization (RI) and directed immobilization (DI), along with that of soluble, surface-free LacZ as control, under both neutral (pH 7) and acidic (pH 4) conditions.

**Figure 4.**
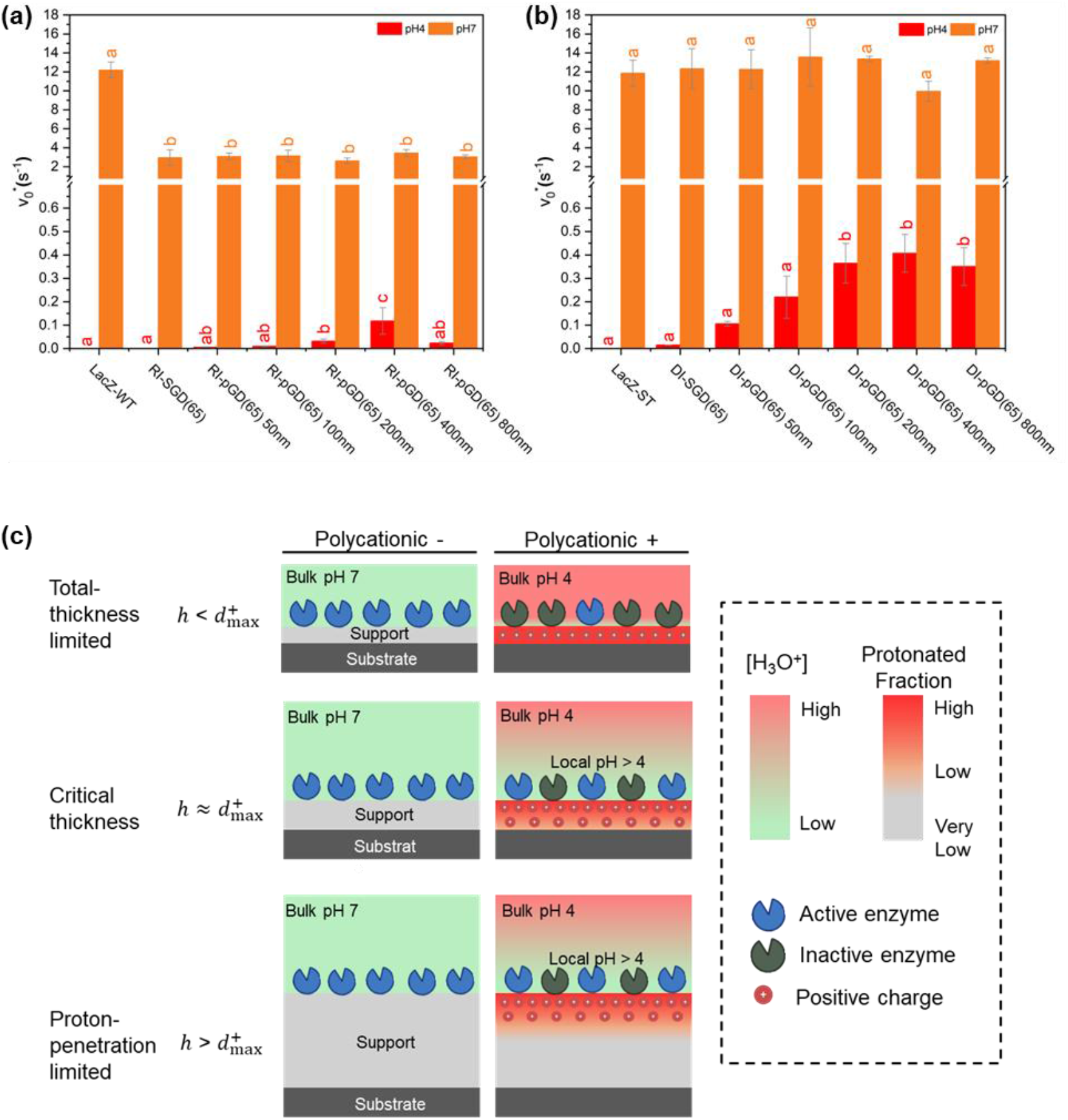
Effect of polymer support thickness on LacZ activity: (a) Normalized initial rate 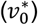 of free and randomly immobilized LacZ-WT (RI) on SGD-modified surface and polymer films with different thickness, at pH 4 and pH 7. (b) Normalized initial rate 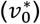 of free and directed immobilized LacZ-ST (DI) under the same conditions. (c) Two-regime conceptual model defined by the relationship between total film thickness, *h*, and the maximum proton penetration depth, 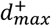: protonation creates a positively charged microenvironment that maintains enzyme function by improving local pH compatibility at the polymer-enzyme interface.

Under neutral conditions (pH 7), randomly immobilized LacZ retained only ∼25% of the catalytic activity of the free enzyme, suggesting that random covalent attachment to surface epoxides imposed conformational constraints and steric hindrance, thereby limiting substrate accessibility.^45^ However, when LacZ was immobilized site-specifically through the DI, the activity was significantly preserved, closely matching that of free enzyme controls. This result underscores the advantage of DI over RI, highlighting that spatial orientation control substantially enhances enzyme function post-immobilization by reducing structural perturbation. Notably, across a support-thickness range of ∼1 to 800 nm, immobilized LacZ activity did not differ significantly (p > 0.05) for either DI or RI. Because protonation is negligible at pH 7, these neutral-pH data serve as a thickness-control, indicating that properties that may covary with total thickness (e.g., mechanical compliance) exert minimal influence on immobilized LacZ activity.

Under acidic conditions (pH 4), the free LacZ showed no detectable catalytic activity, consistent with prior reports that *E. coli* LacZ exhibits maximal activity near neutral pH and strongly reduced activity and stability under acidic conditions.^46^ Remarkably, LacZ immobilized even on the SGD(65) monolayer exhibited measurable catalytic activity, indicating that a single molecular layer of tertiary amines can confer electrostatic proton shielding and/or local pH buffering around the enzyme. Enzyme activity increased monotonically with support thickness from ∼1 nm (SGD(65)) to 200 nm (pGD(65)), with a more pronounced trend for site-directed immobilization (DI) than for random immobilization (RI), reflecting greater activity retention in DI. Beyond a polymer thickness of ∼200 nm (up to 800 nm), no further increase in activity was observed at pH 4 for either immobilization strategy. This plateau aligns with the protonation-depth limit determined earlier (Section 3.2), where polymer layers thicker than ∼250 nm could not undergo additional protonation, indicating a finite proton-penetration depth. Consequently, the activity plateau arises from the saturation of polymer protonation rather than polymer thickness itself—once this limit is reached, additional film thickness offers negligible improvement in local buffering or enzyme protection.

**Figure 4c** summarizes a two-regime conceptual model defined by the relationship between the total thickness of the polycationic support, *h*, and the maximum protonation depth, 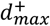. When the film is thinner than the protonation depth 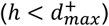, protons can fully penetrate the support and convert tertiary amines into positively charged quaternary amines, placing the system in a **total-thickness-limited regime** (top row). In this regime, very thin supports (e.g., SGD) provide weak proton shielding, whereas increasing *h* increases the amount of protonated polycationic material and correspondingly enhances LacZ activity under acidic stress.

As the film thickness approaches the protonation limit (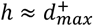 ; middle row), the entire support becomes protonated. Further increases in *h* no longer increase the protonated thickness, defining an optimal or “sweet-spot” thickness that maximizes enzyme protection while minimizing coating material usage and deposition time.

When the support thickness exceeds the protonation depth 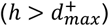; bottom row), the system enters a **proton-penetration-limited regime**, in which the protonated layer thickness remains constant despite additional film growth. Consequently, further increases in total thickness do not improve the acid resistance of immobilized LacZ.

Collectively, these results explicitly demonstrate that the protonation depth, rather than the absolute polymer thickness, determines the extent to which local buffering can protect and preserve enzyme activity under acidic conditions. Controlling polymer thickness within the protonation saturation limit (around 200–300 nm) emerges as a critical strategy for optimizing enzyme immobilization systems for acidic biocatalytic applications.

### 3.4 Influence of Polycationic Monomer density on Immobilized Enzyme Activity

Beyond polymer thickness, the molar ratio of protonatable monomers within polymer formulations could influence the catalytic activity of immobilized enzymes. To systematically investigate this effect, we synthesized copolymers with different compositions of DMAEMA monomer, specifically pGD(25) and pGD(65), representing lower and higher polycationic monomer ratio. Enzyme activities were measured across polymer films of varying thicknesses (200 nm, 400 nm, 800 nm), and results are summarized in **Figure 5**.

**Figure 5.**
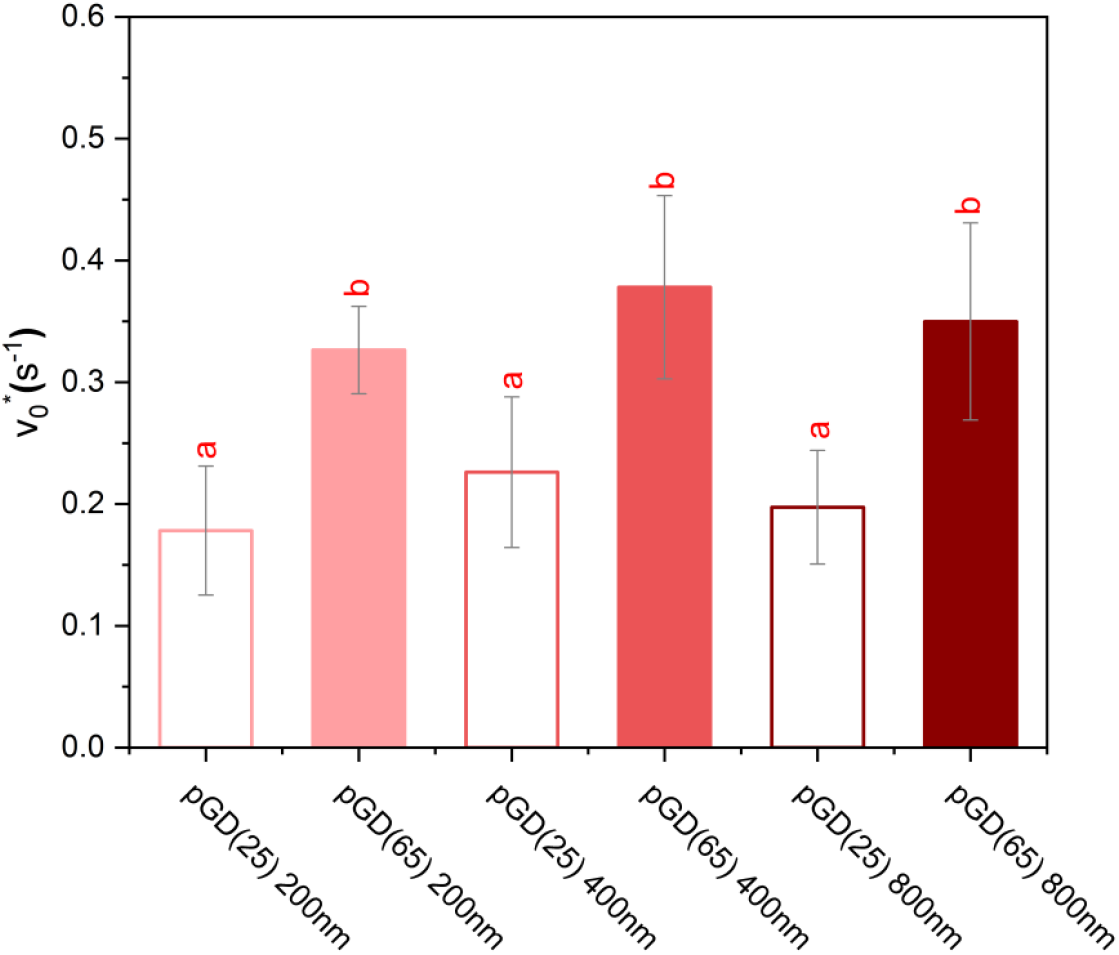
Normalized activities of immobilized LacZ on polymer films with varying thicknesses and compositions, evaluated at pH 4.

Figure 5. shows that for a given polymer composition (either pGD(25) or pGD(65)), no statistically significant difference in enzyme activity was observed among films of different thicknesses. This outcome is consistent with previous observations that, beyond a certain thickness (where protonation saturation occurs), further increases in polymer thickness alone do not influence enzyme activity. However, when we compare enzyme activities immobilized on polymers with different polycationic compositions, a pronounced difference emerged. Specifically, enzyme activity on pGD(65) was significantly higher, 83% greater—than on pGD(25) at all tested thicknesses. This notable increase can be attributed to the substantially higher density of protonatable tertiary amine groups present in pGD(65), providing a stronger local buffering capacity. The underlying mechanism for this protective effect is based on the protonation of tertiary amines under acidic conditions, as indicated by our earlier FTIR results (Section 3.1). Protonation results in positively charged tertiary ammonium groups, which electrostatically repel surrounding hydronium ions (H_3_O_+_) and protons (H_+_), pushing them away from the vicinity of enzyme’s active sites. Nearly all tertiary amines (∼99.99%) in DMAEMA-based polycationic polymer (pKa ∼8.4) are protonatable at pH 4, notwithstanding the aforementioned self-limiting protonation depth behavior. Therefore, the increased DMAEMA content in pGD(65) generates a higher local positive charge density, offering more effective electrostatic repulsion and stronger local buffering capability. Consequently, enzyme immobilized on pGD(65) experienced substantially less acidic stress compared to those on lower-DMAEMA-content polymers such as pGD(25). Our findings align well with previous studies reporting similar protective effects conferred by positively charged polymer supports (e.g., polyethylenimine and chitosan), which have demonstrated significant enzyme activity retention or even enhancements upon immobilization under non-optimal pH conditions.^47,48^

## Conclusion

This study provides a framework for understanding the effect of protonation depth and charge density on polycationic polymer support, and in turn, on enzyme activity under acidic conditions. By combining iCVD and SAM immobilization supports with precisely controlled thickness and tunable copolymer composition, we systematically investigated how film thickness and the molar ratio of protonatable monomers (DMAEMA) influence the spatial distribution and extent of protonation.

We demonstrated that at pH 4 protonation in pGMA-*co*-DMAEMA (DMAEMA∼65%mol) films is confined to a finite nanoscale depth of ∼250 nm, which we term 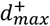, a material parameter that dictates the upper limit of enzymatic protection under the tested conditions. A two-regime model can be defined based on 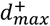: when protonatable film thickness is smaller than 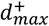, the system is in a total-thickness-limited regime; when protonatable film thickness is greater than 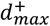, the system is placed in a proton-penetration-limited regime. By correlating this depth with the activity of immobilized LacZ, we showed that matching film thickness to this saturation limit is essential for efficient design: 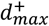 defines an optimal or “sweet-spot” thickness that maximizes enzyme protection while minimizing coating material usage and deposition time. Furthermore, while site-directed immobilization offers better absolute performance by preserving enzyme orientation, the protective buffering mechanism is eventually governed by the polymer’s charge density and protonation depth. These findings shed light on protonation depth and charge density as physicochemical design parameters for developing functional polycationic immobilization supports for enzyme-based biocatalysis, biosensors, and biomedical systems.

## Acknowledgement

Y.C. acknowledges the support of the Foundation for Food and Agriculture Research (FFAR) New Innovator Award (23-000576) and award No. USDA NIFA 2021-67034-35040 for this work. W.S. acknowledges support from start-up funds provided by Virginia Tech. J.C. acknowledges support from the Virginia Tech Biochemistry Graduate Program. The authors thank members of the Cheng and Sun laboratories for technical assistance and constructive feedback.

## Notes

### Competing Interest Statement

The authors have declared no competing interest.

