## Supporting Information for "Nanoscale Protonation Limits and Charge Density in Polymer Films Govern the Activity of Immobilized LacZ under Acid Stress"

**Table S1**. Initiated chemical vapor deposition (iCVD) process parameters for synthesizing the enzyme immobilization supports.

| Samples | Flow rates (sccm) | | | | |
|---|---|---|---|---|---|
| | (Co)monomers | | TBPO | Argon | Total |
| | G | D | | | |
| pGD(25) | 2.0 | 0.6 | 0.6 | 0.0 | 3.2 |
| pGD(65) | 1.0 | 1.0 | 0.6 | 0.4 | 3.0 |

**Table S2**. Pressure saturation ratios (Pm/Psat) of (co)monomer gases and the resulted depostion rates of polymer thin films

| Sample | $P_m/P_{sat}$ (on iCVD stage) | | | Deposition rates (nm $min^{-1}$)* | | |
|---|---|---|---|---|---|---|
| | G | D | Total | $DR_{stg}$ | $DR_{plt}$ | $DR_{stg}/DR_{plt}$ |
| pGD(25) | 0.07 | 0.07 | 0.14 | 96.5 | 5.7 | 16.8 |
| pGD(65) | 0.04 | 0.13 | 0.17 | 46.1 | 0.5 | 94.1 |

*$DR_{stg}$ and $DR_{plt}$ represent the deposition rates measured on a Si substrate placed on the iCVD stage and those measured on another Si substrate located atop a 96-well plate, respectively. The 96-well plates were placed on custom-made aluminum plates that fit the shape of the plates well in order to facilitate heat transfer to the iCVD stage.

**Table S2.** Copolymer molar compositions determined based on transmission FTIR.

| Samples | Copolymer composition (mol%) | |
|---|---|---|
| | **G** | **D** |
| pGD(25) | 76 | 24 |
| pGD(65) | 35 | 65 |

**Table S4.** Primer information

| Primer | Sequences (5' to 3') |
|---|---|
| LacZ-WT-Nde I-F | aagaaggagatatacatatgaccatgattacggattcactggc |
| LacZ-WT-6xHis-R | caaaacagccaagcttttagtgatggtgatggtgatgtttttgacaccagacca act |
| SpyTag-GSlinker-F | tttaagaaggagatatacatatggcgcatatcgtgatggtggatgc |
| SpyTag-GSlinker-R | ttccggatccacctccggatcctttggtcggtttatacgcatccaccatcacgat atg |
| pBAD-LacZ-WT-spy-F | gaggtggatccggaagtggtggatccggaagtggatccggaagtatgacca tgattacgg |
| pBAD-lacz-wt-spy-r | gtatatctccttcttaaagttaaacaaaattatttctagcccaaaaaacgggtat gg |
| SpyCatcher-NdeI-F | aagaaggagatatacatatggttgataccttatcaggtttatcaagtgagc |
| SpyCatcher-HindIII-6xHis-R | caaaacagccaagcttttagtgatggtgatggtgatgtaaatatgagcgtcac ctttag |

### SUPPORTING INFORMATION

#### Gene Blocks information

SpyCatcher:

5’atggttgataccttatcaggtttatcaagtgagcaaggtcagtccggtgatatgacaattgaagaagatagtgct

acccatattaaattctcaaaacgtgatgaggacggcaaagagttagctggtgcaactatggagttgcgtgattcat

ctggtaaaactattagtacatggatttcagatggacaagtgaaagatttctacctgtatccaggaaaatatacattt

gtcgaaaccgcagcaccagacggttatgaggtagcaactgctattacctttacagttaatgagcaaggtcaggtt

actgtaaatggcaaagcaactaaaggtgacgctcatattta3’
